# SLAMF7 engagement super-activates macrophages in acute and chronic inflammation

**DOI:** 10.1101/2020.11.05.368647

**Authors:** Daimon P. Simmons, Hung N. Nguyen, Emma Gomez-Rivas, Yunju Jeong, Antonia F. Chen, Jeffrey K. Lange, George S. Dyer, Philip Blazar, Brandon E. Earp, Accelerating Medicines Partnership (AMP) RA/SLE Network, Deepak A. Rao, Edy Y. Kim, Michael B. Brenner

## Abstract

Macrophages regulate protective immune responses to infectious microbes, but aberrant macrophage activation frequently drives pathological inflammation. To identify regulators of vigorous macrophage activation, we analyzed RNA-seq data from synovial macrophages and identified SLAMF7 as a receptor associated with a super-activated macrophage state in rheumatoid arthritis. We implicated IFN-γ as a key regulator of SLAMF7 expression. Engaging this receptor drove an exuberant wave of inflammatory cytokine expression, and induction of TNF-α following SLAMF7 engagement amplified inflammation through an autocrine signaling loop. We observed SLAMF7-induced gene programs not only in macrophages from rheumatoid arthritis patients, but in gut macrophages from active Crohn’s disease patients and lung macrophages from severe COVID-19 patients. This suggests a central role for SLAMF7 in macrophage super-activation with broad implications in pathology.

## Introduction

Macrophages are necessary for protection against infectious microbes (*1*) but can drive acute inflammation that can become exuberant or chronic and cause significant tissue pathology (*2*). Dysfunctional macrophage activation is evident in autoimmune diseases including rheumatoid arthritis (RA) (*3*–*5*), inflammatory bowel disease (IBD) (*6*–*8*) and interstitial lung disease (*9*, *10*). RA is characterized by infiltration of macrophages into the synovium, along with populations of lymphocytes and activated stromal cells (*11*–*13*). Macrophage numbers and activation states change and can correlate with response to therapy in RA (*14*–*16*). In respiratory infection, macrophages promote inflammation resulting in lung injury (*17*) and contribute to immune activation in severe acute respiratory disease syndrome (*18*, *19*) associated with coronavirus disease of 2019 (COVID-19) (*20*, *21*). Macrophage activation states are determined by receptors for an array of environmental signals (*22*), with cytokines and microbial molecules as the best known macrophage regulators (*23*–*25*). Interferon (IFN)-γ is a key component of classical M1 macrophage activation (*25*) and potentiates macrophage responses to subsequent stimulation (*26*–*28*). Toll-like receptor (TLR) agonists prime macrophages to express inflammasome components that, when activated, result in pyroptotic cell death and release of bioactive interleukin (IL)-1β (*29*). Here, we find that an important component of the macrophage response to a primary signal is upregulation of a secondary super-activator receptor that can then transform these primed or potentiated macrophages into an explosive, potentially pathogenic inflammatory state. As a strategy to find key regulators of macrophage super-activation, we evaluated inflammatory human diseases where macrophages are implicated as major drivers of inflammation. Using this approach, we identified SLAMF7 (also known as CD319, CRACC, and CS1) (*30*–*32*), and implicate this receptor as having a central role in highly activated macrophage-related inflammatory diseases.

## Results

### Marked upregulation of SLAMF7 on macrophages from inflamed synovial tissue

We focused on the inflammatory human disease RA to identify signaling receptors that could act as super-activators of macrophages. We analyzed bulk RNA-sequencing (RNA-seq) data from the Accelerating Medicines Partnership Rheumatoid Arthritis / Systemic Lupus Erythematosus (AMP RA/SLE) Network phase 1 data (*33*) to examine gene expression on sorted CD14+ synovial tissue macrophages from patients with inflammatory RA (n=11) and relatively non-inflammatory osteoarthritis (OA, n=10). Using DESeq2, we identified 509 differentially expressed genes (log_2_foldchange (LFC) ≥ 1, Wald adjusted p value (padj) ≤ 0.05) that were upregulated in RA and that we defined as an “Inflamed RA Macrophage Signature” (Data S1). This signature contained IFN-inducible genes such as *GBP1*, *IFI6*, and *CD40*, as well as inflammatory cytokines and chemokines including *TNF*, *CCL3*, and *CXCL8*, suggesting that both IFN-induced and inflammatory transcriptional programs define the dominant macrophage state in RA. Strikingly, the single most significantly upregulated gene in the “Inflamed RA Macrophage Signature” was *SLAMF7* (LFC=3.82, padj=2.23E-21, Fig. 1A), a receptor that regulates leukocyte activation through homotypic interactions with SLAMF7 on other cells (*30*– *32*). To confirm this finding, we disaggregated synovial tissue from an independent cohort of individuals with OA (n=8, 75% female, mean age 67.6, range 58-85) and RA (n=9, 100% female, mean age 67.7, range 36-86) to quantify SLAMF7 protein expression by flow cytometry (Fig. S1). SLAMF7 was present at very low levels on synovial macrophages from patients with OA but at significantly higher levels on macrophages from patients with RA (Fig. 1B, Mann-Whitney p = 0.0016). We detected SLAMF7 on up to 55 percent of macrophages (mean 26.4%, 95% confidence interval (CI) 13.3-39.53) from patients with RA compared to less than 6 percent of macrophages (mean 2.4, 95% CI 0.35-4.5) from patients with OA (Fig. 1C). We were also able to obtain discarded synovial fluid from de-identified patients with RA and OA. SLAMF7 was expressed at twice the level on synovial fluid macrophages from patients with RA compared to OA (Fig. 1D) and on up to 48 percent of macrophages (mean 24.3, 95% CI 17.03-31.56) from patients with RA, compared to less than 25 percent of macrophages (mean 13.3, 95% CI 6.1-20.5) from patients with OA (Fig. 1E). In contrast, we did not observe major differences in levels of another macrophage-expressed SLAM family member, CD84 (SLAMF5) (*34*) on macrophages from synovial tissue (Fig. S2A-B) or synovial fluid (Fig. S2C-D) from patients with RA versus OA. This associates SLAMF7 as a receptor that is selectively expressed by inflammatory macrophages in RA.

**Fig. 1:**
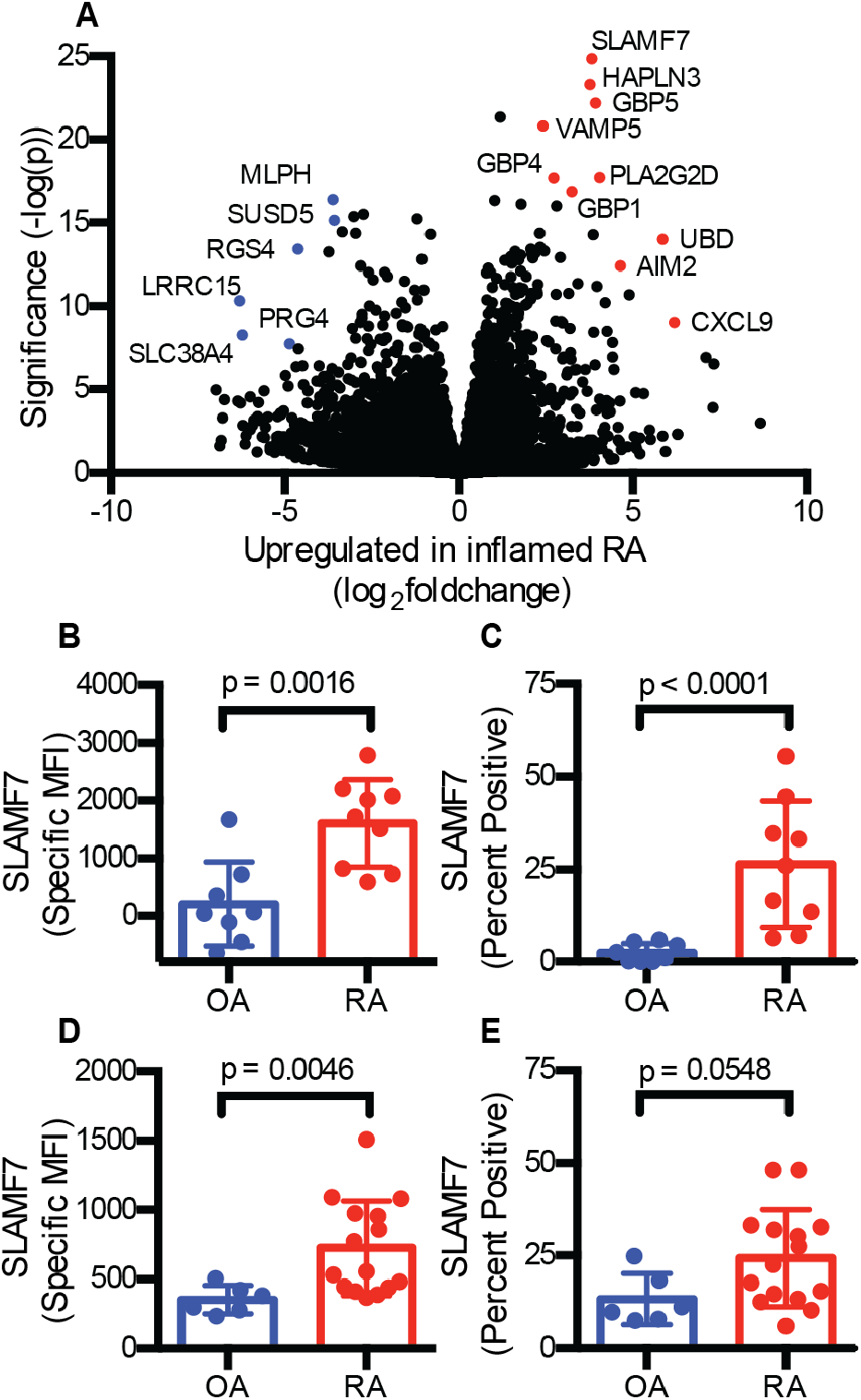
Marked upregulation of SLAMF7 on macrophages from inflamed synovial tissue. A) Differential gene expression in bulk RNA-seq of synovial tissue macrophages from patients with inflamed RA (n=11) compared to OA (n=10). B) Specific MFI for SLAMF7 and C) Percent of macrophages expressing SLAMF7 in synovial tissue from patients with OA (n=8) or RA (n=9). D) Specific MFI for SLAMF7 and E) Percent of macrophages expressing SLAMF7 in synovial fluid from patients with OA (n=6) or RA (n=15). Data represent mean ± SD. The Mann-Whitney test was used for statistical comparisons.

### IFN-γ is a dominant driver of high macrophage SLAMF7 expression

In the unstimulated state, SLAMF7 is expressed at high levels by plasma cells (*35*), and can also be expressed on B cells, T cells, and natural killer (NK) cells, but is expressed at low levels on resting macrophages (*34*, *36*). However, elevated SLAMF7 expression has been found on macrophages from atherosclerotic lesions (*37*) and from patients with myelofibrosis (*38*). BLIMP-1 regulates SLAMF7 expression in lymphocytes (*39*), but it is not well understood how this receptor is regulated in macrophages. To address this gap in knowledge, we sought to use RNA-seq to define the transcriptional activation state of SLAMF7-high macrophages by sorting CD14+ cells with high and low expression of SLAMF7 from peripheral blood of healthy controls (n=5) or patients with RA (n=7), as well as synovial fluid from patients with RA (n=4) (Fig. S3). We then examined genes that were differentially expressed in cells with high versus low SLAMF7 and identified 21 genes that were commonly upregulated (LFC ≥ 1, padj ≤ 0.05) in CD14+ populations of cells with high SLAMF7 expression from both blood and synovial fluid that we defined as a “SLAMF7-High Macrophage Signature” (Data S2). In addition to *SLAMF7*, IFN-inducible genes *CXCL9*, *CD40* and *IDO1* were upregulated in SLAMF7-high CD14+ cells from blood (Fig. 2A) and synovial fluid (Fig. S4A). Gene set enrichment analysis revealed that high SLAMF7 expression was significantly associated (padj <0.05) with msigdb Hallmark gene sets for type I and type II IFN response as well as oxidative phosphorylation (Fig. 2B). IFN signatures were enriched in CD14+ cells with high SLAMF7 from blood and synovial fluid independent of CD16 expression (Fig. S4B), suggesting a prominent *in vivo* role for IFN in regulation of this receptor on macrophages. Oxidative phosphorylation pathways, which have been associated with anti-inflammatory macrophages rather than the glycolytic M1 state (*40*), were enriched in cells with high SLAMF7, suggesting that they are in a poised but not yet fully activated state.

**Figure 2:**
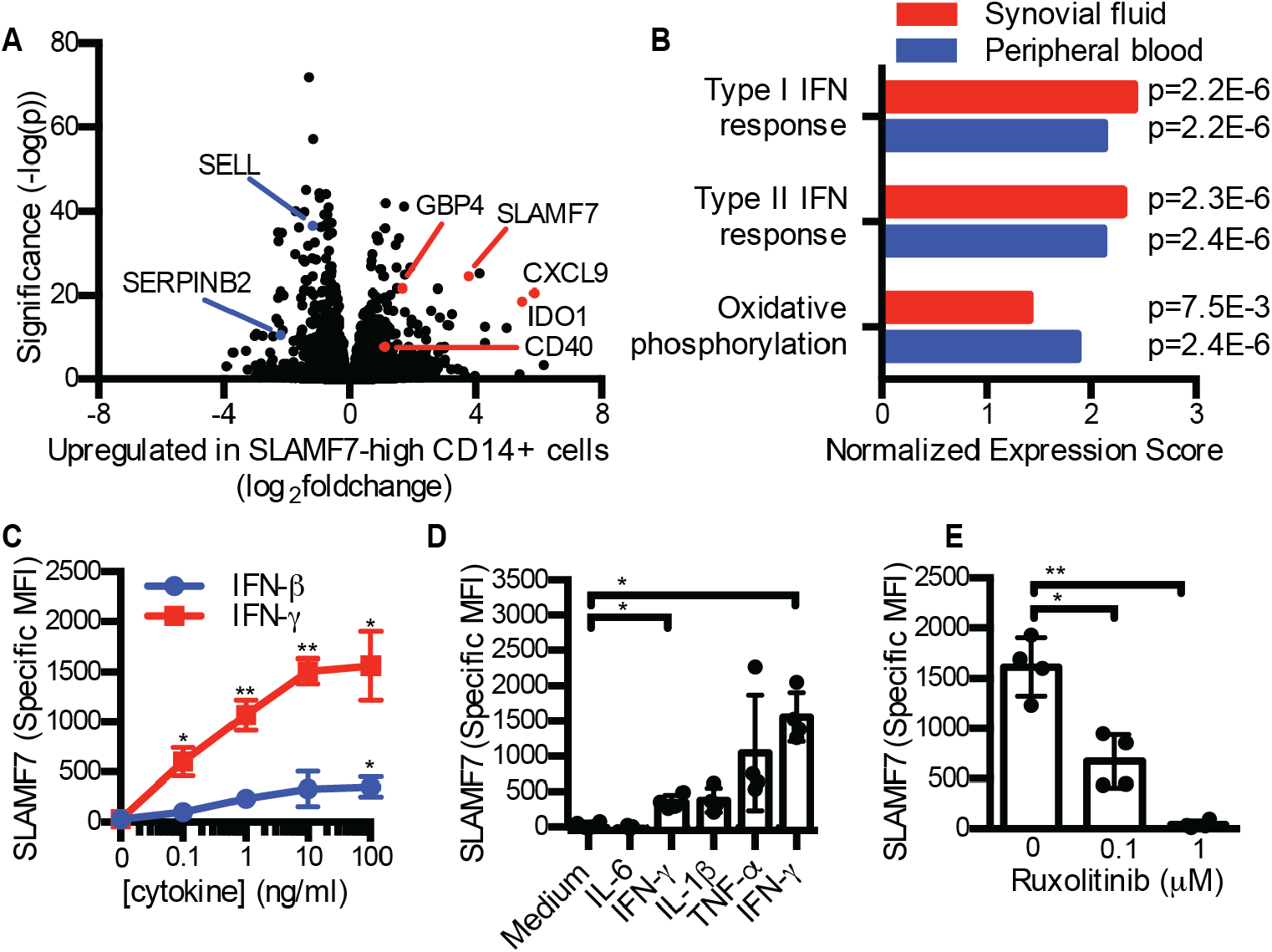
IFN-γ is a dominant regulator of macrophage SLAMF7 expression. A) Differential gene expression in SLAMF7-high versus SLAMF7-low CD14+CD16-cells from peripheral blood (n=12). B) Gene set enrichment analysis for Hallmark Pathways upregulated in SLAMF7-high CD14+CD16-cells from synovial fluid and peripheral blood. C) Specific MFI for SLAMF7 on macrophages incubated with different doses of IFN-γ or IFN-β. D) Specific MFI for SLAMF7 on macrophages incubated with 100 ng/ml of cytokines. IFN-β and IFN-γ results are the same as the 100 ng/ml dose in panel C. E) Macrophages were incubated with ruxolitinib or DMSO prior to IFN-γ treatment (10 ng/ml), and the specific MFI for SLAMF7 was measured after 16h. Data represent mean ± SD of 4 donors. Statistical comparisons were performed using the one-way ANOVA with Dunnett’s multiple comparisons test to compare all cytokines with medium alone. *, p ≤ 0.05; ** p ≤ 0.01.

We performed *in vitro* cytokine stimulation on monocyte-derived macrophages from peripheral blood to distinguish how SLAMF7 is regulated on macrophages. Treatment of macrophages with IFN-γ resulted in high expression of SLAMF7 (Fig. 2C). IFN-β, IL-1β and TNF-α also increased expression of SLAMF7 on macrophages but at lower levels than did IFN-γ, and IL-6 did not affect SLAMF7 expression (Fig. 2D). In contrast, CD84 levels decreased by up to fifty percent after stimulation with IFN-β, IL-1β, TNF-α, and IFN-γ (Fig. S5A-B), while CD45 levels were relatively unchanged (Fig. S5C-D). JAK1 and JAK2 are known to transduce IFN-γ signaling (*41*). Inhibitors of JAK1 and JAK2 such as ruxolitinib effectively disrupt this pathway and are used clinically to treat patients with myelofibrosis (*42*) and graft versus host disease (*43*). We observed profound inhibition of IFN-γ induced SLAMF7 expression in macrophages treated with ruxolitinib *in vitro* (Fig. 2E), confirming the importance of this pathway and revealing a potential method to disrupt SLAMF7 upregulation. Conversely, there was a doubling in CD84 expression in macrophages treated with ruxolitinib (Fig. S5E), suggesting reciprocal regulation of these two SLAM family members by IFN-γ. CD45 expression was unchanged by ruxolitinib treatment (Fig. S5F). Thus, IFN-γ is a dominant regulator of SLAMF7 expression in macrophages.

### Engagement of SLAMF7 triggers an inflammatory cascade

Next, we asked if SLAMF7 might play a role in macrophage activation. SLAMF7-high macrophages from blood and synovial fluid expressed higher levels of IFN-induced genes but not the inflammatory genes from the “Inflamed RA Macrophage Signature.” Based on the striking upregulation of SLAMF7 after primary IFN-γ stimulation, we hypothesized that SLAMF7 engagement might provide a special activating signal for macrophages primed to upregulate its expression. To address this question, we used the 162.1 SLAMF7 activating monoclonal antibody (mAb) (*30*) or recombinant SLAMF7 protein (reported to drive proliferation of myeloma cells (*44*)) to engage SLAMF7 *in vitro* in a two-step process. First, we pretreated macrophages with IFN-γ to induce high SLAMF7 expression, and second we used the activating mAb (a-SLAMF7) or recombinant SLAMF7 protein (r-SLAMF7) to engage cellular SLAMF7. We then performed RNA-seq on these *in vitro* stimulated macrophages to examine transcriptome-wide changes in gene expression. We identified drastic changes in gene expression after SLAMF7 engagement, resulting in 596 upregulated genes driven by both a-SLAMF7 and r-SLAMF7 (LFC ≥ 1, padj ≤ 0.05) that we defined as the “Macrophage SLAMF7 Stimulation Signature” (Data S3). We observed striking upregulation of inflammatory cytokines *TNF*, *IL1B*, *IL6*, and *IL12B* as well as chemokines *CCL3*, *CCL4*, *CXCL1*, *CXCL2*, and *CXCL8* after treatment with activating anti-SLAMF7 mAb (Fig. 3A) or r-SLAMF7 protein (Fig. S6A) compared to macrophages treated with IFN-γ alone. *SLAMF7* was itself upregulated after stimulation, suggesting a positive feedback loop. Note that a large number of cytokines were upregulated in macrophages stimulated with either anti-SLAMF7 or r-SLAMF7, as visualized by heatmap (Fig. 3B). Secreted TNF-α levels increased from 12 pg/ml to 2.2 - 3.0 ng/ml after stimulation (Fig. 3C) and IL-6 levels increased from 6 pg/ml to 0.7 - 1.2 ng/ml after stimulation (Fig. 3D). Real-time PCR (RT-PCR) analysis confirmed induction of *CCL3* (mean LFC 3.1 - 4.2, Fig. S6B), *CXCL1* (mean LFC 4.3 - 5.1, Fig. S6C), and *CXCL8* (mean LFC 10.0 - 10.2, Fig. S6D) after SLAMF7 engagement.

**Figure 3.**
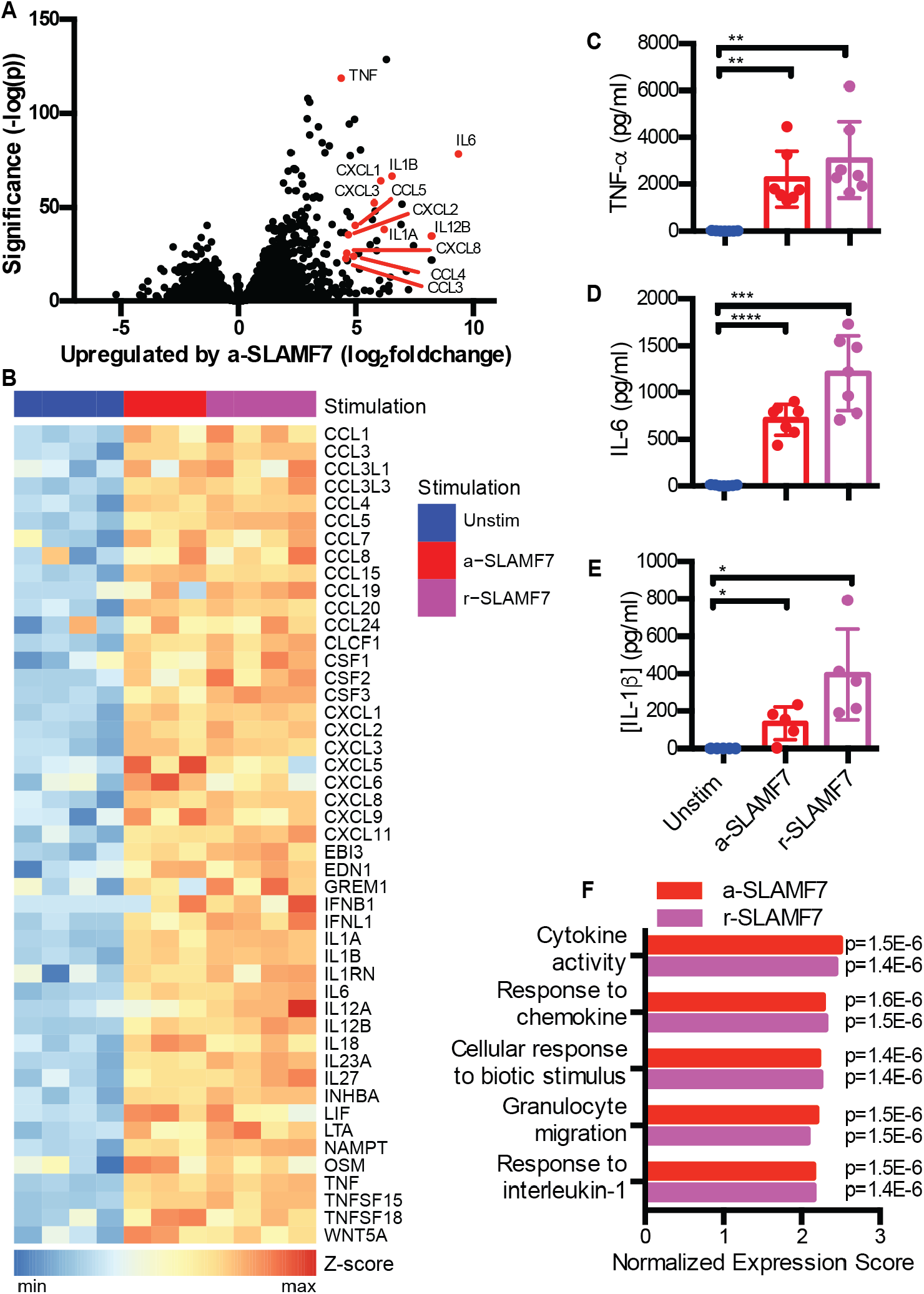
Engagement of SLAMF7 triggers an inflammatory cascade. Macrophages were potentiated with IFN-γ (10 ng/ml) for 24 hours prior to treatment with a-SLAMF7 (10 μg/ml) or r-SLAMF7 (1 μg/ml) for 4h. A) Differential gene expression for macrophages incubated with a-SLAMF7 (n=3 donors) compared to macrophages treated with only IFN-γ (n=4 donors). B) Heatmap showing z-scores for gene expression values of differentially expressed genes in the msigdb GO:cytokine activity gene set for IFN-γ pre-treated macrophages without additional stimulation (n=4 donors), stimulated with a-SLAMF7 (n=3 donors), or with r-SLAMF7 (n=4 donors). C) Secreted TNF-α, and D) secreted IL-6 after macrophage incubation with a-SLAMF7 or r-SLAMF7 (n=7 donors). E) Release of IL-1β after macrophage incubation with a-SLAMF7 or r-SLAMF7 for 4h followed by treatment with nigericin (10 μM) for 30m (n=5 donors). Data represent mean ± SD. F) Gene set enrichment analysis for macrophages after stimulation with a-SLAMF7 or r-SLAMF7 showing enrichment for GO:0005125, cytokine activity; GO:1990868, response to chemokine; GO:0071216, cellular response to biotic stimulus; GO:0097530, granulocyte migration; and GO:0070555, response to interleukin-1. Statistics were calculated with the one-way ANOVA, using Dunnett’s multiple comparisons test to compare each condition to medium alone. *, p ≤ 0.05; **, p ≤ 0.01; ***, p ≤ 0.001; ****, p ≤ 0.0001; unstim, unstimulated; a-SLAMF7, anti-SLAMF7 antibody; r-SLAMF7, recombinant SLAMF7 protein.

The inflammatory cytokine *IL1B* was strongly upregulated after stimulation with SLAMF7. However, macrophages only express high levels of inflammasome components and pro-IL-1β when they are primed by microbial TLR agonists or cytokines like TNF-α, and subsequent inflammasome activation with pyroptotic cell death is required for abrupt release of bioactive IL-1β (*29*, *45*). We detected release of 134 - 395 pg/ml of IL-1β from macrophages stimulated with SLAMF7 when nigericin was added to activate the inflammasome (Fig. 3E), suggesting that SLAMF7 can also prime inflammasomes. Gene set enrichment analysis identified strong enrichment of msigdb gene ontology (GO) categories (padj <0.05) including cytokine activity, cellular response to biotic stimulus, and myeloid leukocyte migration (Fig. 3F), indicating that triggering SLAMF7 drives a dominant myeloid inflammatory program. We did not observe a strong overlap with the msigdb Immunologic Signatures gene set for classically activated M1 macrophages. The “Classical M1 vs. Alternative M2 Macrophage” upregulated gene set (*46*) defines activation after 18h combined stimulation with IFN-γ + LPS, but interestingly the array of cytokines driven by SLAMF7 engagement was strikingly absent from that signature. Other M1 activation protocols include initial priming with IFN-γ followed by LPS, which results in production of inflammatory cytokines and gene expression (*47*) that partially overlaps with those we identified in our conditions. This underscores the fact that this SLAMF7 activation program rests upon and is a separate step after primary stimulation of macrophages by IFN-γ or other M1 differentiation and activation factors. These sequential *in vitro* conditions include 1) initial macrophage potentiation with IFN-γ to drive high SLAMF7 expression, followed by 2) engagement of SLAMF7 and completion of activation, underscoring this distinct program. We termed this activation state, defined by upregulation of the SLAMF7 receptor followed by SLAMF7 engagement that then triggers profound inflammatory activation, as the super-activated macrophage inflammatory state induced by SLAMF7 engagement (SAM7).

### SLAMF7 amplifies macrophage activation through a TNF-α autocrine loop

Macrophage activation by SLAMF7 engagement was so striking that we wondered if it might recruit autocrine amplification pathways. A time course revealed the extremely rapid induction of TNF-α within 2 hours, with continued accumulation over time (Fig. 4A). Based on the very rapid induction of TNF-α, we hypothesized that TNF might provide an autocrine amplification signal as previously noted in other contexts (*26*, *48*). Indeed, antibody blockade of TNF-α decreased expression of *TNF* (Fig. 4B) and *IL1B* (Fig. 4C) by fifty percent after SLAMF7 stimulation. Corresponding blockade of the TNFR1 and TNFR2 receptors also reduced levels of *TNF* and *IL1B* (Fig. 4B-C). We also used siRNA to silence the receptors TNFR1 and TNFR2, resulting in a marked reduction of TNF-α secretion after stimulation with r-SLAMF7 (Fig. 4D). We confirmed the selectivity and effectiveness of siRNA silencing for the genes encoding these receptors, *TNFRSF1A* (Fig. S7A) and *TNFRSF1B* (Fig. S7B), by RT-PCR. This implicates TNF- α autocrine signaling as an additional amplification step for inflammatory pathway activation following SLAMF7 engagement in SAM7 macrophages.

**Figure 4.**
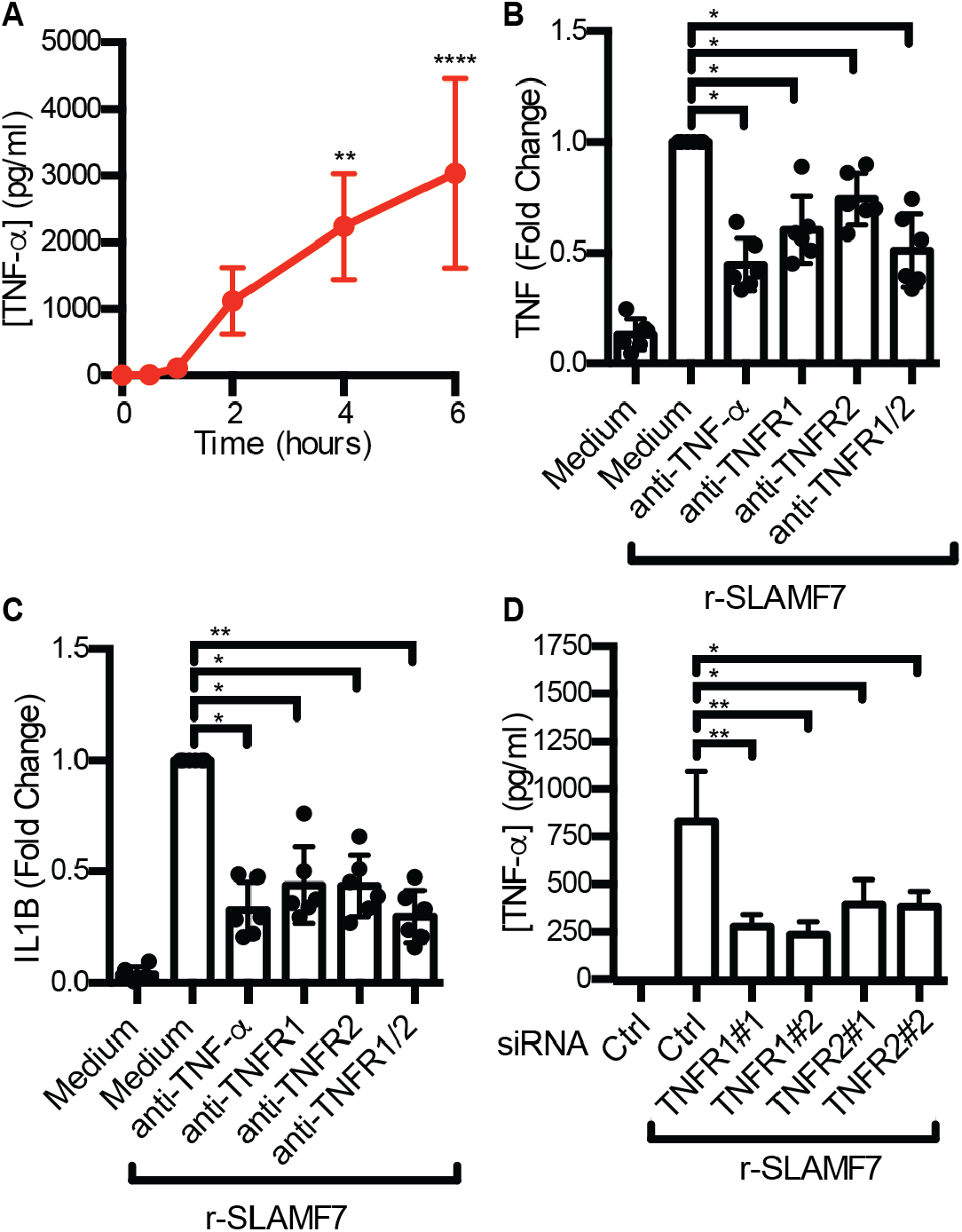
SLAMF7 amplifies macrophage activation through a TNF-α autocrine loop. A) Macrophages were potentiated with IFN-γ (10 ng/ml) for 24h, then stimulated with r-SLAMF7 (500 ng/ml). Secreted TNF-α was measured by ELISA. Data represent mean ± SD of four donors. B-C) Macrophages were potentiated with IFN-γ (10 ng/ml) for 24h, treated with antibodies for 30m, and stimulated with r-SLAMF7 (100 ng/ml) for 8h. RT-PCR was used to quantify B) *TNF* and C) *IL1B* relative to macrophages without antibody pre-treatment. Data represent mean ± SD of 6 donors. D) Macrophages were treated with control siRNA or two different siRNAs targeting TNFR1 or TNFR2, potentiated with IFN-γ (5 ng/ml) for 16-18h, and stimulated with r-SLAMF7 (100 ng/ml) for 3h. TNF-α was measured by ELISA. Data represent mean ± SD of triplicate wells from an experiment representative of 3 independent experiments. Statistics were calculated using the one-way ANOVA with Dunnett’s multiple comparisons test. *, p ≤ 0.05; **, p ≤ 0.01; ****, p ≤ 0.0001; r-SLAMF7, recombinant SLAMF7 protein.

### SLAMF7 super-activation of macrophages in autoimmune and infectious disease

To determine whether SAM7 macrophages contribute broadly to disease pathology, we correlated our *in vitro* findings with molecular data from diseased human tissues. As described above, we developed two distinct SLAMF7 macrophage signatures: an *in vivo* signature from sorted cells with high SLAMF7 expression, and another from cells stimulated *in vitro* by this receptor. We first analyzed the differentially expressed genes we had identified from macrophages in inflamed RA from AMP RA/SLE Network Phase 1 data (*33*) using the 21 genes in the “SLAMF7-High Macrophage Signature” (derived from sorted cells with high SLAMF7 expression), and we identified 11 genes that were significantly upregulated (LFC ≥ 1, padj ≤ 0.05) and none that were significantly downregulated (LFC ≤ 1, padj ≤ 0.05) in inflamed RA. Then, using the *in vitro* derived “Macrophage SLAMF7 Stimulation Signature”, we found that 121 genes were significantly upregulated while only 12 out of the 596 genes in this signature were significantly downregulated in inflamed RA (Fig. 5A). We used gene set enrichment analysis to quantify this overlap and determined that both signatures were highly enriched in the synovial macrophage gene expression program from patients with RA (Fig. 5B). We then derived a “SLAMF7 Activation Score” defined as the percent of total gene expression derived from the 221 most highly upregulated genes (LFC ≥ 2, padj ≤ 0.01) in the “Macrophage SLAMF7 Stimulation Signature.” The average SLAMF7 activation score was almost twice as high in macrophages from individuals with RA than OA, suggesting that this program strongly contributes to the inflammatory gene expression program in RA (Fig. 5C).

**Figure 5:**
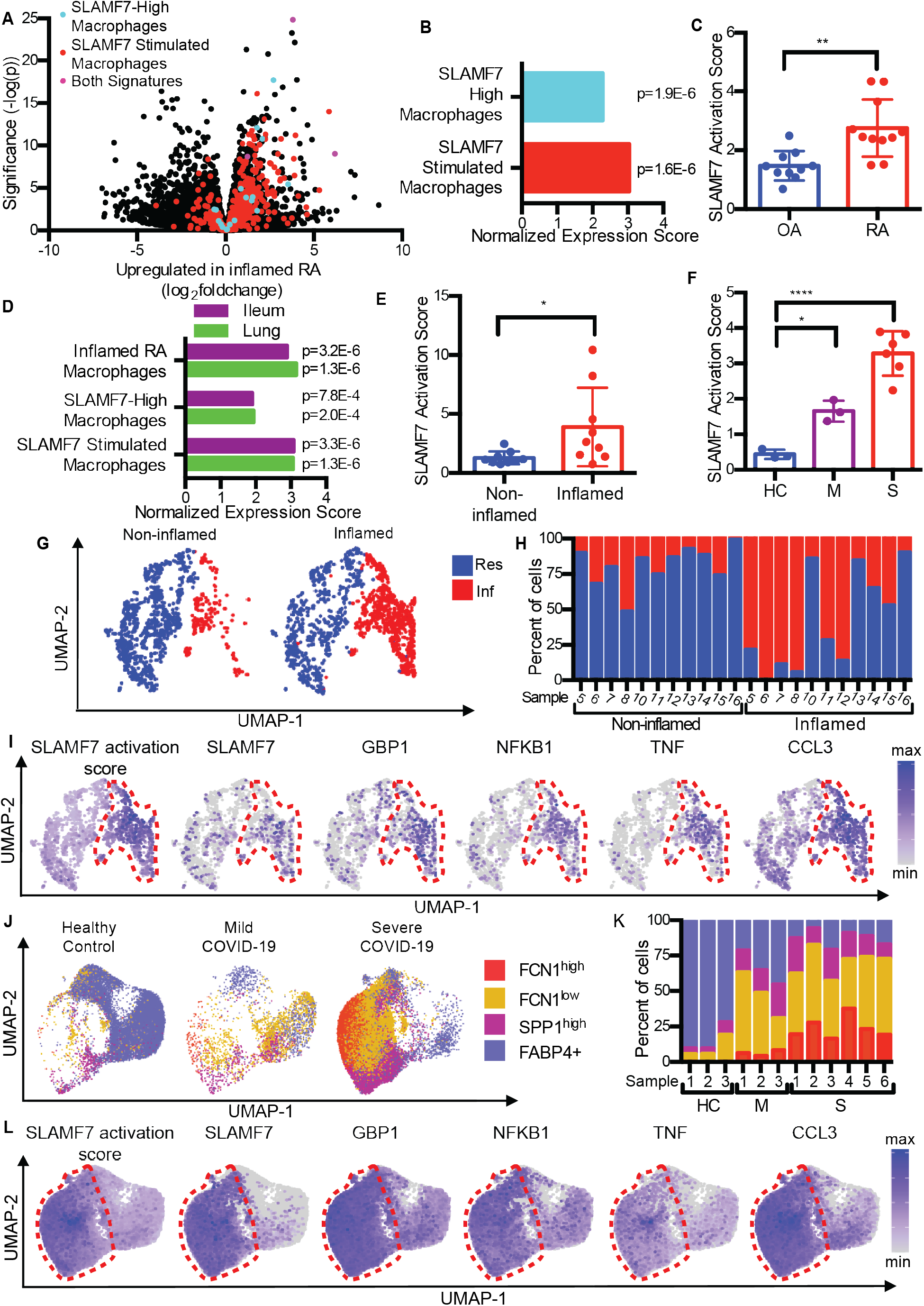
SLAMF7 super-activated macrophages drive inflammation in autoimmune and infectious disease. A) Volcano plot from Fig. 1A highlighting genes from the “SLAMF7-High Macrophage Signature,” the “Macrophage SLAMF7 Stimulation Signature,” and genes included in both signatures. B) Gene set enrichment analysis comparing differential gene expression in RA versus OA to the “SLAMF7-High Macrophage Signature” and “Macrophage SLAMF7 Stimulation Signature.” C) SLAMF7 activation score for bulk RNA-seq data on synovial macrophages from patients with OA (n=10) or RA (n=11). Data represent mean ± SD. D) Gene set enrichment analysis comparing gene expression from macrophages from inflamed ileal tissues in patients with Crohn’s disease or lungs of patients with COVID-19 with the “Inflamed RA Macrophage Signature”, the “SLAMF7-High Macrophage Signature” and the “Macrophage SLAMF7 Stimulation Signature.” E) SLAMF7 activation score for macrophages from non-inflamed (n=9) and inflamed ileal tissues (n=9). F) SLAMF7 activation score for bronchoalveolar lavage macrophages from healthy controls (n=3), or individuals with mild (n=3) or severe COVID-19 (n=6). Data in E-F represent mean ± SD. G) UMAP plot of macrophage clusters from involved and uninvolved ileal tissues. H) Percent of macrophages from each donor assigned to each cluster. I) UMAP plots showing gene expression of ileal macrophage populations. J) UMAP plot of bronchoalveolar lavage macrophage populations. K) Percent of macrophages from each donor assigned to each population. L) UMAP plots showing gene expression for bronchoalveolar lavage macrophage populations. The paired t-test was used for two-way statistical comparisons, and the one-way ANOVA with Dunnett’s multiple comparisons test was used to compare mild and severe COVID-19 to healthy controls.*, p ≤ 0.05; **, p ≤ 0.01; ****, p < 0.0001; Res, Resident macrophage cluster; Inf, Inflammatory macrophage cluster; HC, healthy control; M, mild COVID-19; S, severe COVID-19.

Macrophages contribute to inflammatory pathways in Crohn’s disease (*6*–*8*) as part of the GIMATS (IgG plasma cells, inflammatory mononuclear phagocytes, and activated T and stromal cells) module that has been associated with resistance to TNF-blockade in patients with IBD (*8*), and macrophages also are major contributors to pathological inflammation in individuals with COVID-19 pneumonia (*18*, *19*). We analyzed publicly available single cell RNA-seq data on ileal tissue from patients with Crohn’s disease (*8*) and bronchoalveolar lavage fluid from individuals infected with COVID-19 (*18*) to determine whether this SLAMF7 activation program identified in RA contributes to macrophage-driven inflammation in other diseases. First, we used gene set enrichment analysis to compare the gene expression profiles of macrophages from inflamed ileal tissue and infected lungs to genes upregulated in RA that we had defined as the “Inflamed RA Macrophage Signature.” We observed significant overlap of macrophage gene expression in both IBD and COVID-19 with gene expression in RA (Fig. 5D), suggesting common pathways of macrophage activation. We then compared the transcriptomes of macrophages from inflamed ileal tissue and bronchoalveolar lavage cells with the *in vivo* derived “SLAMF7-High Macrophage Signature” and the *in vitro* derived “Macrophage SLAMF7 Stimulation Signature.” Importantly, we found a strong correlation between these signatures and gene expression in macrophages from inflamed gut and lung tissues (Fig. 5D). The average levels of SLAMF7 activation were more than twice as high in macrophages from inflamed gut tissue than non-inflamed tissue (Fig. 5E), with the highest levels of SLAMF7 activation in samples categorized as having high (Fig. S8A) rather than low (Fig. S8B) GIMATS module intensity scores. Individuals with mild COVID-19 lung involvement had three times higher levels of SLAMF7 activation than healthy controls, and remarkably, the signature was more than six-fold higher in individuals with severe disease (Fig. 5F). This suggests that this SAM7 gene expression program represents a dominant macrophage state in these inflammatory diseases.

We then explored which single cell populations displayed the SAM7 macrophage super-activation state *in vivo*. First, we visualized macrophage populations from ileal tissue of patients with IBD using Uniform Manifold Approximation and Projection (UMAP) plots. We observed two dominant populations of ileal macrophages with expression of *CD14* and *CD68*, including a population of macrophages with higher expression of *MRC1* (Fig. S9) that was present in both inflamed and non-inflamed tissues, consistent with “resident macrophage” populations (Fig. 5G). Another macrophage population with higher cytokine expression was predominantly derived from inflamed tissues, consistent with “inflammatory macrophages” (Fig. 5G, H). Gene expression from the “SLAMF7 Activation Score” was extremely high in the inflammatory macrophage population (Fig. 5I, red circle). We confirmed that these cells had higher expression of *SLAMF7*, as well as *GBP1*, *NFKB1*, *TNF*, *CCL3* (Fig. 5I, S10), IFN-induced *CD40* and *IDO1*, and inflammatory chemokines *CCL4* and *CXCL8* (Fig. S11).

We then focused on lung macrophages from patients infected with COVID-19, including a monocyte-like *FCN1*^high^ population, a chemokine-high *FCN1*^low^ group, fibrosis-associated *SPP1*^high^ macrophages (*10*), and *FABP4*+ alveolar macrophages (Fig. 5J). There was clear separation of the three *FCN1+* and *SPP1*+ macrophage populations from the *FABP4*+ alveolar macrophages (Fig. S12), with drastic increases in the proportions of *FCN1*^high^, *FCN1*^low^, and *SPP1*^high^ macrophage populations in individuals with COVID-19 (Fig. 5K). We observed the highest levels of the “SLAMF7 Activation Score” in cells from the *FCN1^l^*^ow^ and *FCN1*^high^ populations that were expanded in patients with severe disease, although it was highly expressed in all three *FCN1*+ and *SPP1*+ populations compared to *FAB4*+ alveolar macrophages (Fig. 5L, red circle). There was also a striking increase in *SLAMF7* expression in macrophages from patients with severe COVID-19 compared to extremely low expression in healthy controls (Fig. 5L, S13). These populations also expressed very high levels of *GBP1*, *NFKB1*, *TNF* and *CCL3* (Fig. 5L, S13), as well as IFN-inducible *CD40*, *IDO1*, and inflammatory *CCL4* and *CXCL8* (Fig. S14). The convergence of higher levels of *SLAMF7* and other IFN-induced genes, along with SLAMF7-induced inflammatory genes, may implicate the SAM7 state as an important contributor to COVID-19 pneumonia. This macrophage activation state, defined by initial potentiation with IFN-γ and upregulation of *SLAMF7*, followed by subsequent SLAMF7 engagement and amplification through TNF-α, is one that may play a major role in pathological inflammation of RA, IBD and COVID-19 pneumonia.

## Discussion

Macrophages orchestrate immune responses to protect against invading microbes, but overly robust immune responses and inflammation can cause autoimmune disease or cytokine storm during infection. Here, we have identified a pathway in which SLAMF7 is a key receptor that is highly upregulated on macrophages exposed to IFN-γ, with lesser but significant induction by other cytokines. Subsequent triggering of SLAMF7, even in the absence of microbial molecules, drives rapid and explosive production of cytokines and chemokines that we have defined as a super-activated macrophage inflammatory state induced by SLAMF7 engagement (SAM7). SLAMF7 has been reported to inhibit cytokine production by monocytes (*49*, *50*) but also to activate phagocytosis by murine macrophages (*51*) under different conditions than what we have used here. The sequential process defined here with IFN-γ stimulation followed by SLAMF7 engagement results in a strong inflammatory program. Because macrophages must pass through multiple steps to achieve this complete super-activation state, interventions targeting different parts of this pathway could allow for fine-tuned therapeutic strategies. JAK inhibitors such as ruxolitinib impair macrophage responses to IFN-γ, and would diminish the initial upregulation of SLAMF7, thereby limiting activation of this pathway. TNF-α blockade could impair autocrine amplification of this pathway but would have limited effects on the array of inflammatory effectors driven by primary SLAMF7 stimulation. However, it is blockade of SLAMF7 itself that would prevent the completion of macrophage super-activation on IFN-γ potentiated macrophages and likely have the most important therapeutic implications. The absence of this receptor on normal resident macrophage populations implicates it as having particular importance as a specific target for inflammatory macrophages. Elotuzumab is an antibody targeting SLAMF7 that is in clinical use for treatment of relapsed multiple myeloma (*52*), and part of this drug’s efficacy is through activation of NK cells (*53*) and macrophages (*54*). Another antibody against SLAMF7 was used to deplete plasma cells and reduce disease severity in an animal model of arthritis (*55*), but the importance of this receptor on highly activated macrophages has not previously been appreciated. SLAMF7 and its activation signature are strikingly upregulated in inflammatory RA, IBD, and COVID-19 pneumonia, implicating it as contributing to highly activated macrophage-driven inflammation in autoimmune and infectious diseases. Targeting this receptor may offer a special opportunity to block severe macrophage-driven inflammation without diminishing more moderate macrophage immune surveillance and helpful homeostatic macrophage functions.

## Materials and Methods

### Human research

Human subjects research was performed with approval from the Institutional Review Board at Brigham and Women’s Hospital. Patients with rheumatoid arthritis (RA) were defined according to ACR 2010 Rheumatoid Arthritis classification criteria (*56*). Synovial tissue samples were obtained as excess samples from patients undergoing arthroplasty procedures at Brigham and Women’s Hospital, and samples were frozen in Cryostor CS10 preservation medium (Sigma-Aldrich). Synovial fluid samples were obtained sterile prior to arthrotomy as discarded samples from routine clinical care, and peripheral blood samples were collected as discarded samples from routine clinical care. Patient consent for genomic studies was obtained for all samples used for RNA sequencing.

### Sample processing

Synovial tissue was disaggregated as previously described (*11*, *57*). Frozen synovial tissue was thawed and minced into small pieces that were digested with 100 μg/ml Liberase TL (Sigma) and 100 μg/ml DNAse I (Sigma) at 37°C for 30 minutes. Samples were passed over a 70 μM cell strainer (Fisher), RBCs were lysed with ACK lysis buffer (Fisher), and cells were stained for flow cytometry. Synovial fluid and peripheral blood were isolated as previously reported (*58*), with density centrifugation on a Ficoll-Hypaque gradient to isolate mononuclear cells. Cells were cryopreserved for subsequent analysis.

### Reagents and antibodies

Recombinant human M-CSF, IFN-β, IFN-γ, IL-1β, IL-6 and TNF-α were from Peprotech. ELISA kits for TNF-α, IL-6, and IL-1β were from R&D Systems. Ultra-LEAF purified anti-TNF (MAb1) was from BioLegend. Anti-TNFR1 (16803) and anti-TNFR2 (22210) were from R&D Systems. Ruxolitinib was from Seleckchem. Nigericin was from Invivogen. Fetal bovine serum (FBS) was from Hyclone. Propidium iodide was from Biolegend. Recombinant SLAMF7 protein (r-SLAMF7) was from Sigma. Purified anti-SLAMF7 (162.1) was from BioLegend. The following antibodies were used for flow cytometry and cell sorting: CD16 (CB16, eBioscience), CD45 (HI30, BioLegend), CD14 (61D3, eBioscience), SLAMF7 (162.1, BioLegend), CD84 (CD84.1.21, BioLegend), CD3 (UCHT1, eBioscience), CD19 (HIB19, eBioscience), CD56 (CMSSB, eBioscience), mouse IgG2b, κ isotype control (MPC-11, BioLegend).

### Cell culture and stimulation

Monocytes were MACs purified from peripheral blood mononuclear cells using CD14 microbeads (Miltenyi). 50,000 cells per well were cultured in flat bottom 96 well plates in RPMI (Fisher) with 10% FBS, with additives of beta-mercaptoethanol, sodium pyruvate, HEPES, penicillin/streptomycin, and L-glutamine (Fisher) and M-CSF (20 ng/ml). Macrophages were allowed to rest for 24 h. Cytokines were then added for an additional 24 h. For inhibitor experiments, ruxolitinib or an equivalent concentration of DMSO was added 30 minutes prior to IFN-γ. For SLAMF7 stimulation, after 24 h incubation with IFN-γ, macrophages were treated with anti-SLAMF7 antibody or r-SLAMF7 protein. For blocking experiments, antibodies were added at 10 μg/ml (TNFR1 and TNFR2) or 20 μg/ml (TNF-α) 30 minutes prior to stimulation with r-SLAMF7.

### siRNA

Monocytes were cultured in RPMI with 10% FBS and 20 ng/ml M-CSF for 1 week. They were then transfected with either a control siRNA or siRNA against a gene of interest (Silencer Select, Life Technologies) at 30 nM by reverse transfection using RNAiMax (Life Technologies). The following day, medium was replaced with RPMI containing 1% FBS. After 24 hours of siRNA treatment, macrophages were treated with IFN-γ for 16-24h and stimulated with recombinant SLAMF7 protein.

### Quantitative Real-Time PCR (RT-PCR)

Primers were from IDT. RNA was processed using the RNEasy Mini kit (Qiagen) and Quantitect Reverse Transcription kit (Qiagen). RT-PCR was performed using SybrGreen (Agilent) on the AriaMX (Agilent). Gene expression for each sample was normalized to *GAPDH* and compared across conditions.

### Cell staining

Cells were washed with staining buffer (PBS, 2% FBS, 2mM EDTA) (Fisher), treated with Human Trustain FcX (Biolegend) for 10 minutes on ice, and stained with antibodies for 30 minutes on ice. Propidium iodide (1 μg/ml) was added 15 minutes prior to acquisition. For experiments with stimulated macrophages, cells were stained and then fixed in 4% paraformaldehyde for 10 minutes on ice prior to acquisition. Analysis was done on the LSR-Fortessa (BD).

### Flow cytometry analysis

Analysis was performed by gating on myeloid cells by forward scatter and side scatter, excluding doublets, and gating on CD45+ leukocytes with exclusion of dead cells. CD14+ cells were selected for analysis (Fig. S1). Specific MFI was determined by subtracting the MFI from an isotype control from the MFI for a specific antibody. Percent positive cells were calculated by setting a gate with ≤ 1% of cells in a sample stained with isotype control as the threshold for positivity.

### Cell sorting

Sorting was performed on a FACS-Aria Fusion sorter (BD). Myeloid cells were gated as CD45+ cells with exclusion of live/dead dye, CD3, CD19 and CD56. SLAMF7-low cells were defined as 40% of myeloid cells with lowest expression of SLAMF7. SLAMF7-high cells were defined as those with staining above the level of isotype control. Three populations from blood and two populations from synovial fluid were sorted based on expression of CD14 and CD16 (Fig. S3). Up to 1,000 cells from each population were collected in Eppendorf tubes with 5 μl of TCL buffer and 1% beta-mercaptoethanol. Cell lysates were frozen at −80 for further processing.

### RNA-seq

Cell lysates were collected in TCL buffer with 0.1% beta-mercaptoethanol and frozen at −80°C. Samples were sequenced at the Broad Institute using the Smart-seq2 RNA-seq platform (*59*–*61*). Samples with detection of at least 10,000 genes were used for subsequent analysis.

### Data analysis

Flow cytometry data were analyzed using Flowjo version 10.4 (Treestar). Graphical and statistical analysis was done in RStudio version 1.1.383 with R version 3.6.0, JMP Pro version 13.0.0 (JMP Inc), and Prism version 6.0.h (GraphPad). Kallisto version 0.46.0 was used for quantification of RNA-seq reads (*62*) using version GRCh38.97 of the Ensembl transcriptome (downloaded August 13, 2019). Differential gene expression analysis was performed using DESeq2 version 1.24.0 (*63*). Gene set enrichment analysis was performed with fgsea version 1.10.1 (*64*) using msigdb gene sets (downloaded December 4, 2019), including Hallmark gene sets (h.all.v7.0.symbols.gmt), Gene Ontology gene sets (c5.all.v7.0.symbols.gmt) and Immunologic Signature gene sets (c7.all.v7.0.symbols.gmt). Heatmaps were generated using pheatmap version 1.0.12. Single cell RNA-seq analysis was performed on a cloud-computing cluster using R version 3.5.3, Seurat version 3.1.4 (*65*, *66*), and harmony version 1.0 (*67*).

### Analysis of bulk RNA-seq from synovial tissue macrophages

FASTQ files for bulk RNA-seq data were obtained for sorted CD14+ synovial macrophages from patients with OA (n=10) and inflamed RA (n=11) from Accelerating Medicines Partnership (AMP) RA/SLE Network phase 1 (*33*) (dbGaP phs001457.v1.p1). Reads were quantified using kallisto. To focus on signaling receptors, we selected genes with the “protein_coding” Ensembl biotype with at least 1 count across samples, resulting in 18,304 genes for analysis. DESeq2 was used for differential gene expression analysis comparing macrophages from inflamed RA and OA, including both disease status and processing batch in the model. The 509 genes with log_2_foldchange ≥ 1, adjusted p value ≤ 0.05 for RA compared to OA were considered significantly upregulated genes, defined as the “Inflamed RA Macrophage Signature” (Data S1).

### Analysis of bulk RNA-seq from sorted SLAMF7-high and SLAMF7-low cells

RNA-seq on SLAMF7-high and SLAMF7-low cells sorted from peripheral blood from healthy controls (n=5) and patients with RA (n=7), and synovial fluid from patients with RA (n=4 donors), was performed at the Broad Institute using Smart-seq2 with 25 bp paired reads. Read quantification was performed using the Broad Institute pipeline, with alignment to GRCh38.83 using STAR version 2.4.2a (*68*) and quantification with RSEM version 1.2.2.1 (*69*). 37,414 genes were included for analysis. DESeq2 was used to calculate differential gene expression between SLAMF7-high and SLAMF7-low cells for each population, including both SLAMF7 expression and donors in the model. Genes with log_2_foldchange ≥ 1, adjusted p value ≤ 0.05 were considered significantly upregulated, and the “SLAMF7-High Macrophage Signature” was defined as 21 genes that were upregulated in SLAMF7-high compared to SLAMF7-low CD14+CD16-cells from both peripheral blood and synovial fluid (Data S2). Differentially expressed genes from SLAMF7-high versus SLAMF7-low cells from each population in peripheral blood and synovial fluid were ordered by the Wald statistic calculated by DESeq2. The fgsea package was used for gene set enrichment analysis with 10^6^ permutations for comparison of these ordered genes to msigdb Hallmark gene sets (h.all.v7.0.symbols.gmt).

### Analysis of bulk RNA-seq from *in vitro* SLAMF7 stimulated macrophages

For analysis of macrophages after *in vitro* SLAMF7 stimulation (n=4 donors), RNA-seq was performed using Smart-seq2 with 50 bp paired reads. Reads were quantified using kallisto. One sample stimulated with a-SLAMF7 was excluded due to low gene counts. 36,513 genes were included for analysis. DESeq2 was used for differential gene expression analysis comparing macrophages potentiated with IFN-γ, followed by stimulation with either anti-SLAMF7 or r-SLAMF7 to macrophages treated with IFN-γ without additional stimulation. Fold-change values were shrunk using the apeglm algorithm (*70*). Genes with log_2_foldchange ≥ 1, adjusted p value ≤ 0.05 were considered significantly upregulated. The “Macrophage SLAMF7 Stimulation Score” was defined as 596 genes that were commonly upregulated with both a-SLAMF7 and r-SLAMF7 stimulation (Data S3). Differentially expressed genes from macrophages stimulated with a-SLAMF7 or r-SLAMF7 compared to unstimulated macrophages were ordered by the Wald statistic calculated by DESeq2. The fgsea package was used for gene set enrichment analysis with 10^6^ permutations for msigdb Gene Ontology gene sets (c5.all.v7.0.symbols.gmt).

### Analysis of single cell RNA-seq on macrophages from ileal samples from individuals with Crohn’s disease

Read count matrices for single cell RNA-seq of ileal biopsies from patients with Inflammatory Bowel Disease (IBD) (GSE134809) (*8*) were downloaded on December 10, 2019. Cells were filtered to include those with > 500 genes and < 5,000 genes, and < 25% mitochondrial RNA. Harmony reduction was used to correct for differences between samples for clustering and UMAP analysis. Myeloid cells were selected for analysis based on expression of *CD68* and *CD14*. From the myeloid population, we excluded non-macrophage populations of dendritic cells (DCs) and other cells to focus our analysis on macrophages, using *CD1C* and *FCER1A* to identify CD1c+ DCs and monocyte-derived DCs, *CLEC9A* and *BATF3* to identify CD141+ DCs, *CCR7* and *LAMP3* to identify migratory DCs, *GZMB* and *IRF7* to identify plasmacytoid DCs, and *CD3D* and *IGHG1* to exclude possible doublets. The remaining macrophages were positive for *CD14*, *CD68*, *CSF1R*, and *MAFB*. A cluster of resident macrophages (Res) had higher expression of *MRC1*, *CD163* and *MERTK*, while a cluster of inflammatory macrophages (Inf) had high expression of inflammatory cytokines and chemokines including *TNF* and *CCL3* (*8*). As described in the original publication, samples from donor 6 were excluded from additional analysis because of the low number of cells and samples from donor 16 were excluded due to similarity in cells from involved and uninvolved tissues.

### Analysis of single cell RNA-seq from bronchoalveolar lavage of individuals with COVID-19 infection

Read count matrices for single cell RNA-seq of bronchoalveolar lavage cells from patients with COVID-19 (GSE145926) (*18*) were downloaded on May 5, 2020 and metadata provided by the authors was downloaded from github on May 21, 2020. We selected myeloid cells and populations in the metadata as originally described for analysis (*18*). Cells were filtered to include those with genes > 500 and < 5,000. Harmony reduction was used to correct for differences between donors for UMAP visualization.

### Custom gene-set enrichment analysis

For single cell datasets, the sum of counts for each gene from all cells for each sample was combined to generate pseudobulk RNA-seq counts. For ileal macrophages from GSE134809, DESeq2 was used for differential gene expression comparing inflamed tissues (n=9) compared to non-inflamed tissues (n=9), including both disease involvement and donor in the model. For bronchoalveolar lavage macrophages from GSE145926, DESeq2 was used for differential gene expression comparing healthy controls (n=3) to patients with severe COVID-19 infection (n=6), including disease in the model. Differentially expressed genes for macrophages from inflamed versus non-inflamed tissues from patients with RA or IBD, or healthy versus COVID-19 infected individuals were ordered by the Wald statistic calculated by DESeq2. Genes from the “Inflamed RA Macrophage Signature”, the “SLAMF7-High Macrophage Signature” and “Macrophage SLAMF7 Stimulation Signature” were compiled into a custom derived gmt file. The fgsea package was used with 10^6^ permutations for gene set enrichment analysis of these ordered gene lists against these custom gene sets.

### SLAMF7 activation score

The “SLAMF7 Activation Score” was calculated using the most highly upregulated genes (log_2_foldchange ≥ 2, adjusted p value ≤ 0.01, n=221 genes) from the “Macrophage SLAMF7 Stimulation Signature.” For bulk RNA-seq data on CD14+ synovial macrophages from AMP, the “SLAMF7 Activation Score” was calculated as the sum of counts for these 221 genes as a percent of total gene counts for each donor. For single cell RNA-seq data on macrophages from patients with IBD or COVID-19, the “SLAMF7 Activation Score” was calculated as the sum of counts for these 221 genes as a percent of total gene counts for each cell, and the median value for all cells from each donor was used for analysis.

## Acknowledgments

We thank Fan Zhang, Ilya Korsunsky and Soumya Raychaudhuri for advice on bioinformatic analysis, Mike Gurish and Greg Keras for assistance recruiting patients and processing samples, Bill Apruzzese for helpful discussion, and patients for their participation.

