## Supplemental Figures for "SLAMF7 engagement super-activates macrophages in acute and chronic inflammation"

**Primer List**

| Primer | Sequence |
|---|---|
| GAPDH | Forward: AATCCCATCACCATCTTCCAG<br>Reverse: AAATGAGCCCCAGCCTTC |
| CCL3 | Forward: CGGCAGATTCCACAGAATTTC<br>Reverse: AGGTCGCTGACATATTTCTGG |
| CXCL1 | Forward: AACCGAAGTCATAGCCACAC<br>Reverse: CCTCCCTTCTGGTCAGTTG |
| CXCL8 | Forward: ATACTCCAAACCTTTCCACCC<br>Reverse: TCTGCACCCAGTTTTCCTTG |
| TNF | Forward: ACTTTGGAGTGATCGGCC<br>Reverse: GCTTGAGGGTTTGCTACAAC |
| IL1B | Forward: ATGCACCTGTACGATCACTG<br>Reverse: ACAAAGGACATGGAGAACACC |
| TNFSFR1A | Forward: TGCCAGGAGAAACAGAACAC<br>Reverse: TCCTCAGTGCCCTTAACATTC |
| TNFSFR1B | Forward: GTCCACACGATCCCAACAC<br>Reverse: TGTCACACCCACAATCAGTC |

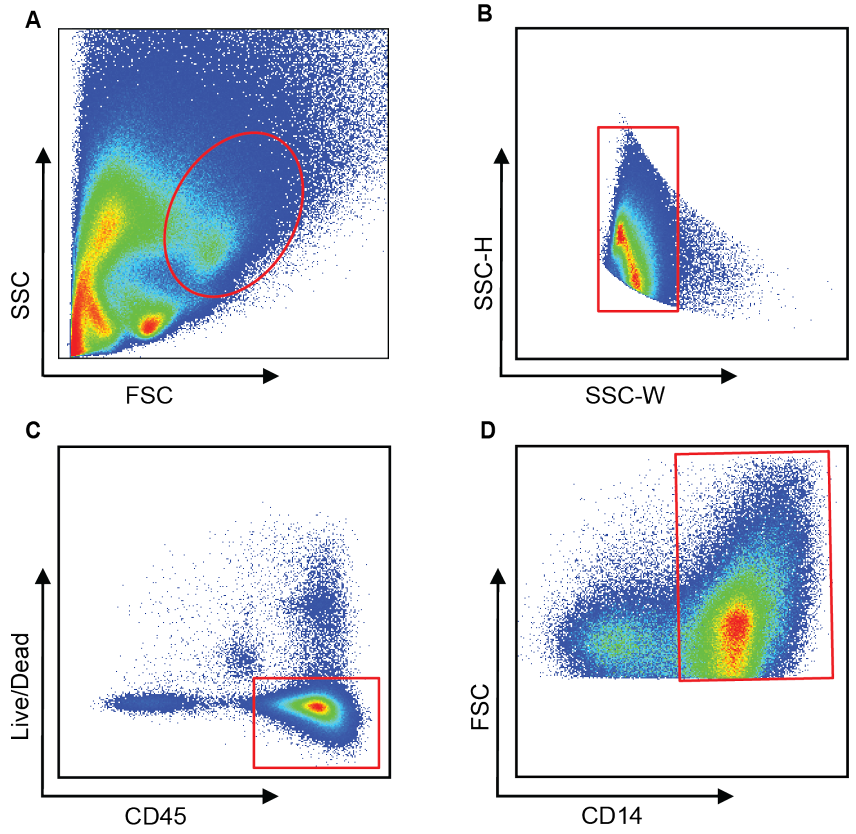


**Fig. S1. Gating for flow cytometric analysis of macrophages.** A) Myeloid cells were selected based on forward and side scatter. B) Doublets were excluded with side scatter height versus side scatter width. C) CD45+ cells were selected with exclusion of live/dead dye. D) CD14+ cells were selected for analysis.

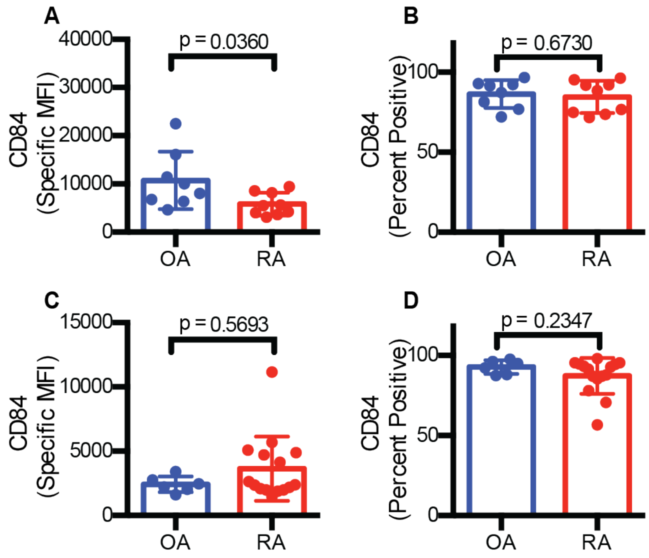


**Fig. S2. CD84 expression in synovial macrophages.** A) Specific MFI for CD84, and B) Percent of macrophages expressing CD84 in synovial tissue from patients with OA (n=8) or RA (n=9). C) Specific MFI for CD84 , and D) Percent of macrophages expressing CD84 in synovial fluid from patients with OA (n=6) or RA (n=15). Data represent mean ± SD. The Mann-Whitney test was used for statistical comparisons.

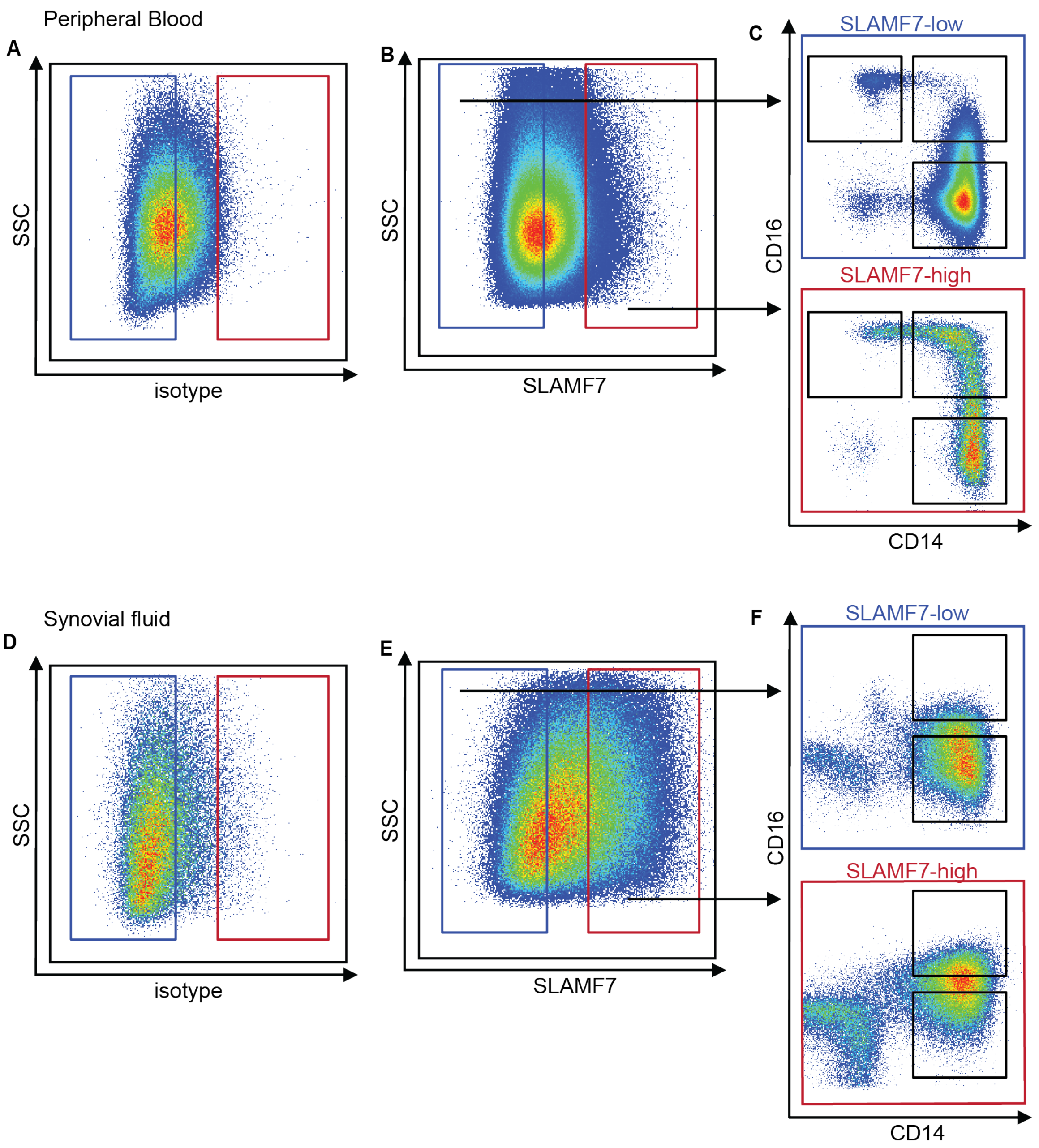


**Fig. S3. Gating strategy for cell sorting.** Myeloid cells were selected by gating on live CD45+ cells as in Fig. S1, with exclusion of lymphocytes by CD3, CD19 and CD56. A) An isotype control was used to define the threshold for SLAMF7-positive cells from peripheral blood. B) Gating to sort SLAMF7-low and SLAMF7-high cells. C) Populations of monocytes defined by CD14 and CD16 expression were sorted from SLAMF7-low (outlined in blue) and SLAMF7-

high (outlined in red) cells from peripheral blood. D) An isotype control was used to define the threshold for SLAMF7-positive cells from synovial fluid. E) Gating to sort SLAMF7-low and SLAMF7-high cells. F) Macrophage populations defined by CD14 and CD16 expression were sorted from SLAMF7-low (outlined in blue) and SLAMF7-high (outlined in red) cells from synovial fluid.

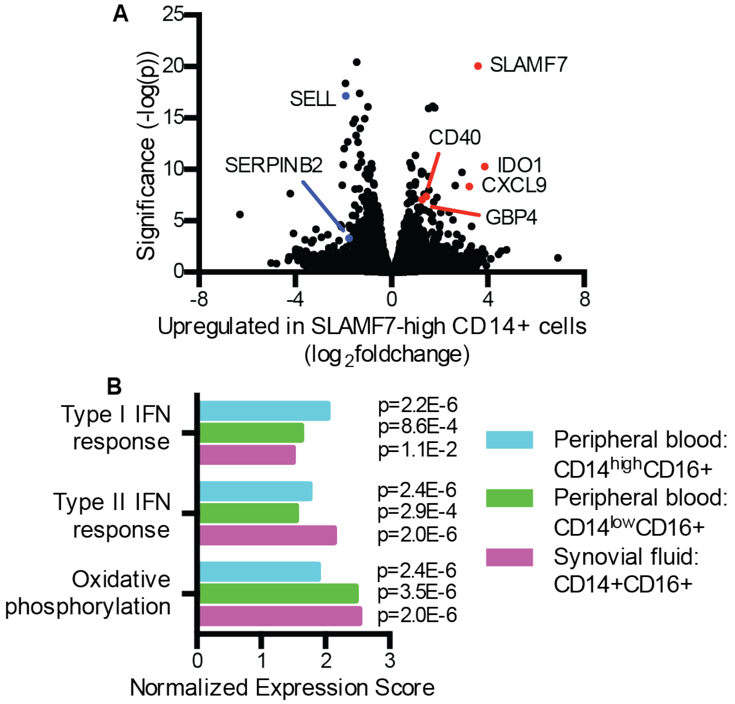


**Fig. S4. SLAMF7 expression *in vivo* associated with interferon signature.** A) Differential gene expression for SLAMF7-high compared to SLAMF7-low CD14+CD16- cells from synovial fluid (n=4 donors). B) Gene set enrichment analysis for Hallmark Pathways upregulated in SLAMF7-high CD14+CD16+ cells from synovial fluid and peripheral blood.

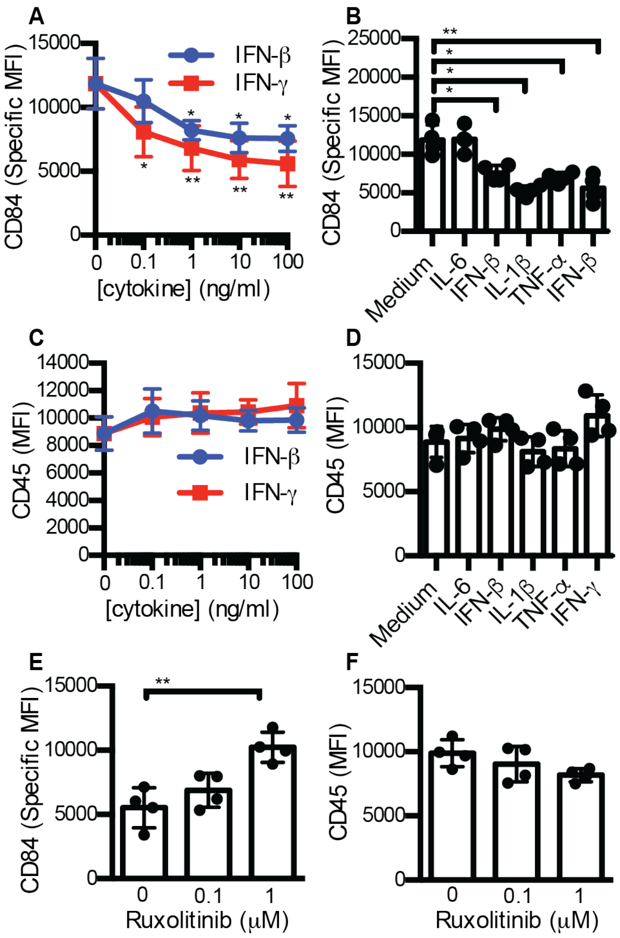


**Fig. S5. Macrophage surface marker expression after cytokine stimulation.** A) Specific MFI for CD84 on macrophages incubated with different doses of IFN-γ or IFN-β. B) Specific MFI for CD84 on macrophages incubated with 100 ng/ml of cytokines. IFN-β and IFN-γ results are the same as the 100 ng/ml dose in panel A. C) MFI for CD45 on macrophages incubated with different doses of IFN-γ or IFN-β. D) MFI for CD45 expression on macrophages incubated with 100 ng/ml of cytokines. IFN-β and IFN-γ results are the same as the 100 ng/ml dose in panel C. E-F) Macrophages were incubated with ruxolitinib or DMSO prior to IFN-γ treatment (10 ng/ml). Specific MFI for E) CD84 and MFI for F) CD45 was measured after 16h. Data represent mean ± SD of 4 donors. Statistical comparisons were performed using the one-way ANOVA with Dunnett's multiple comparisons test to compare all cytokine conditions with medium alone.

*, $p \leq 0.05$; ** $p \leq 0.01$.

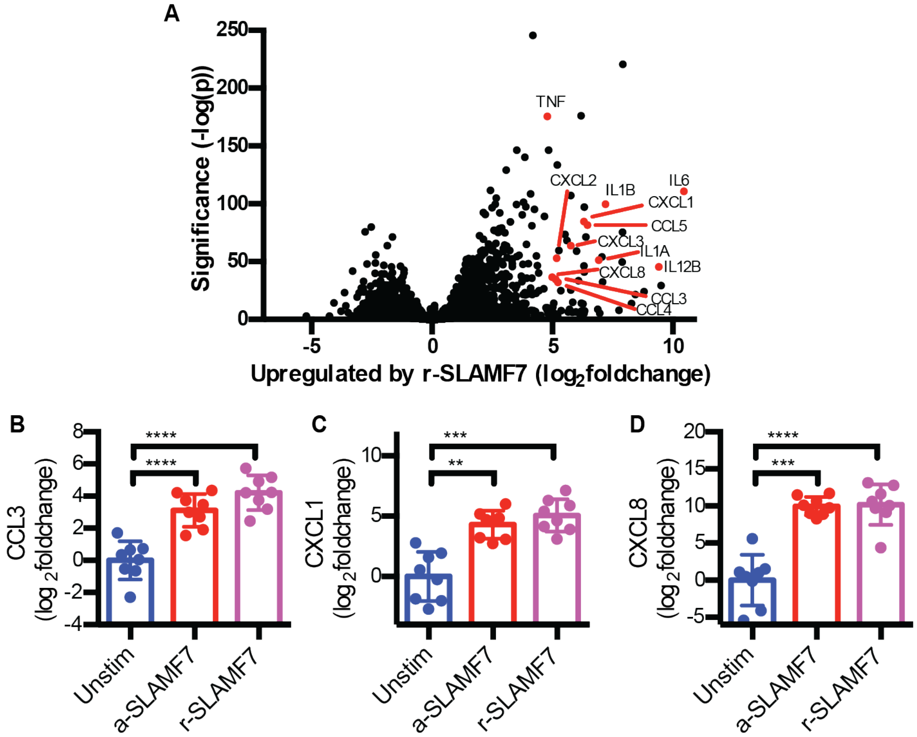


**Figure S6. Inflammatory cascade driven by SLAMF7 engagement.** Macrophages were potentiated with IFN-γ (10 ng/ml) for 24 hours prior to subsequent treatment with a-SLAMF7 (10 µg/ml) or r-SLAMF7 (1 µg/ml) for 4h. A) Differential gene expression for macrophages incubated with r-SLAMF7 for 4h (n=4) compared to unstimulated macrophages (n=4). B-D) RT-PCR was used to quantify expression of B) CCL3, C) CXCL1, and D) CXCL8 relative to unstimulated macrophages. Data represent mean ± SD of 8 donors. The one-way ANOVA with Dunnett's multiple comparisons test was used to compare each condition to medium alone. **, $p \leq 0.01$; ***, $p \leq 0.001$; ****, $p \leq 0.0001$; unstim, unstimulated; a-SLAMF7, anti-SLAMF7 antibody; r-SLAMF7, recombinant SLAMF7 protein.

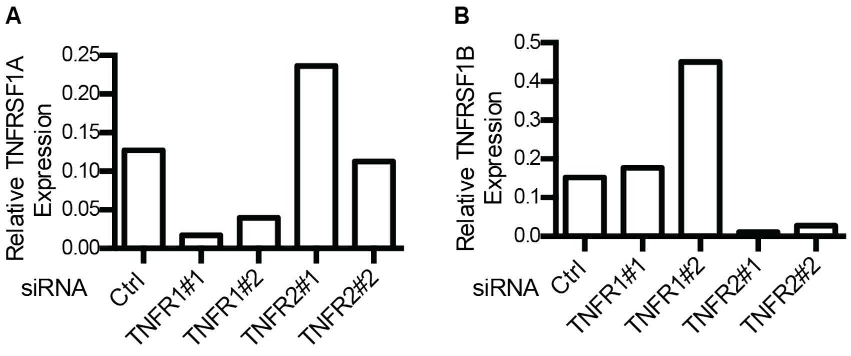


**Figure S7: Confirmation of siRNA silencing.** Gene expression of A) TNFRSF1A and B) TNFRSF1B was quantified in macrophages treated with control siRNA, or two different siRNAs targeting either TNFR1 or TNFR2. Data represent the mean of duplicate samples.

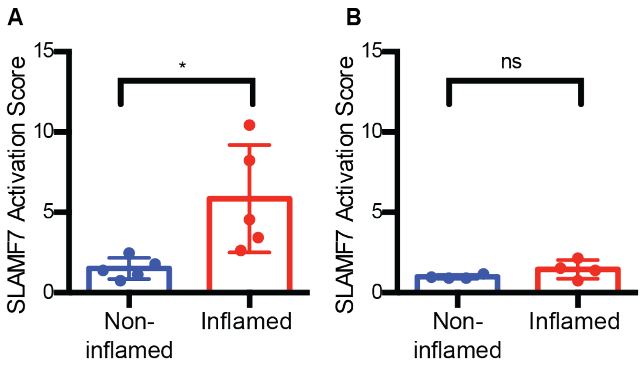


**Figure S8: High SLAMF7 activation in ileal macrophages from samples with high GIMATS module intensity scores.** A) SLAMF7 activation score for donors reported to have a high GIMATS module intensity score (n=5). B) SLAMF7 activation score for donors reported to have a low GIMATS module intensity score (n=4). Data represent mean ± SD. The paired t-test was used for statistical comparisons. *, $p \leq 0.05$.

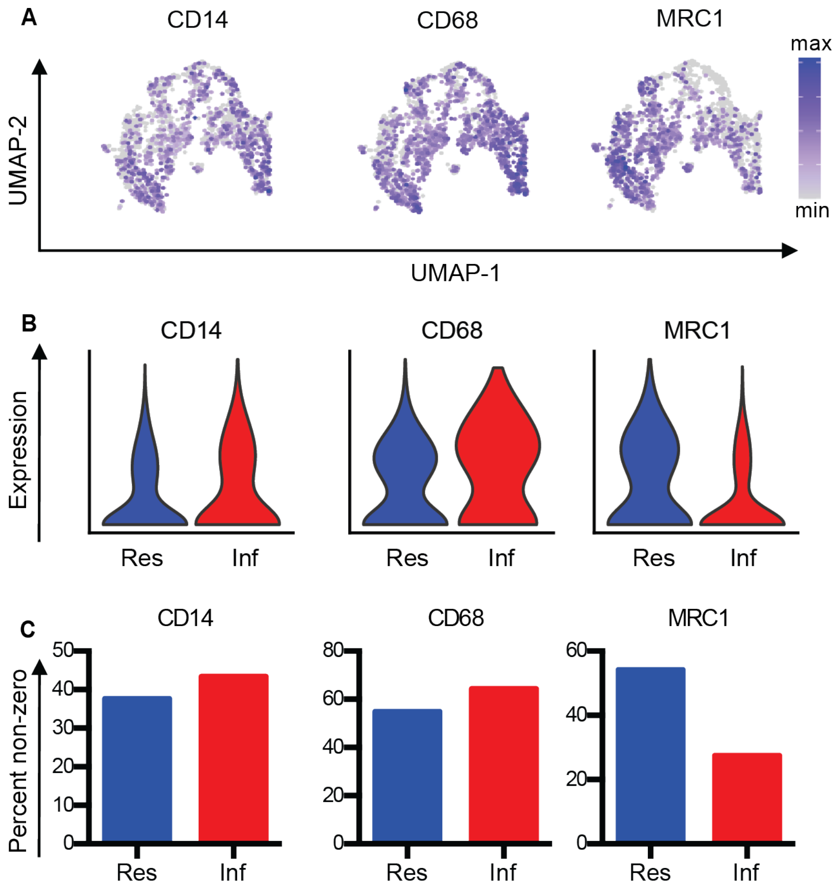


**Figure S9: Macrophage populations in ileal tissue from patients with inflammatory bowel disease.** A) UMAP plot from Fig. 5G showing gene expression values in ileal macrophage populations. B) Violin plots of gene expression in macrophage clusters from Fig. 5G. C) Percent of cells with expression of genes in macrophage clusters from Fig. 5G. Res, resident macrophages; Inf, Inflammatory macrophages.

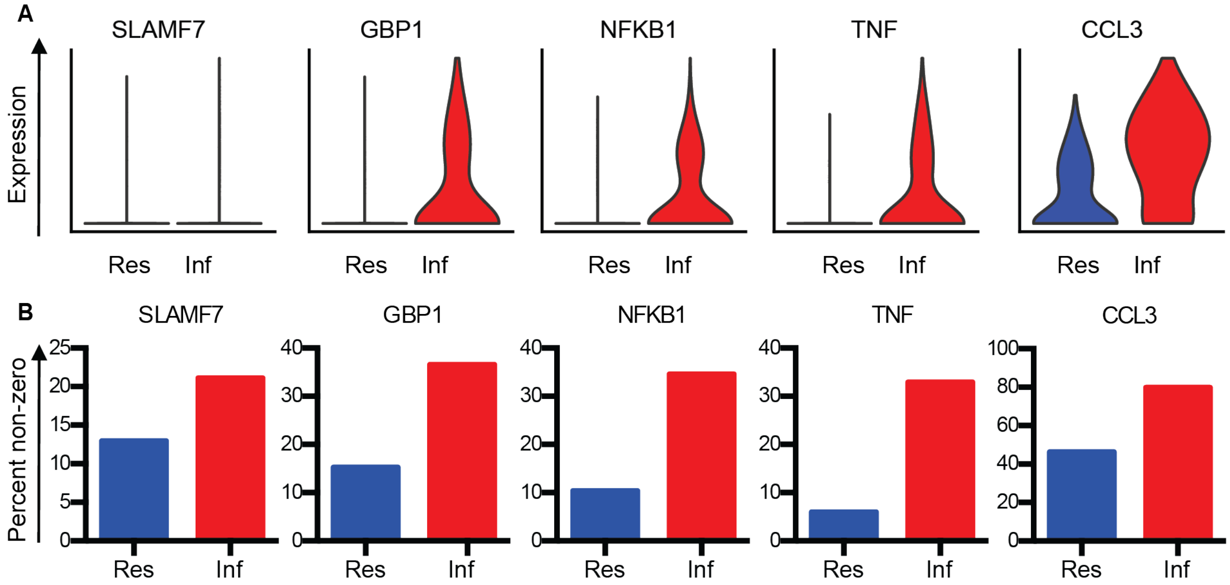


**Figure S10: Evidence for SLAMF7 activation in macrophages from patients with inflammatory bowel disease.** A) Violin plots of gene expression in macrophage clusters from Fig. 5G. B) Percent of cells with expression of genes in macrophage clusters from Fig. 5G. Res, resident macrophages; Inf, Inflammatory macrophages.

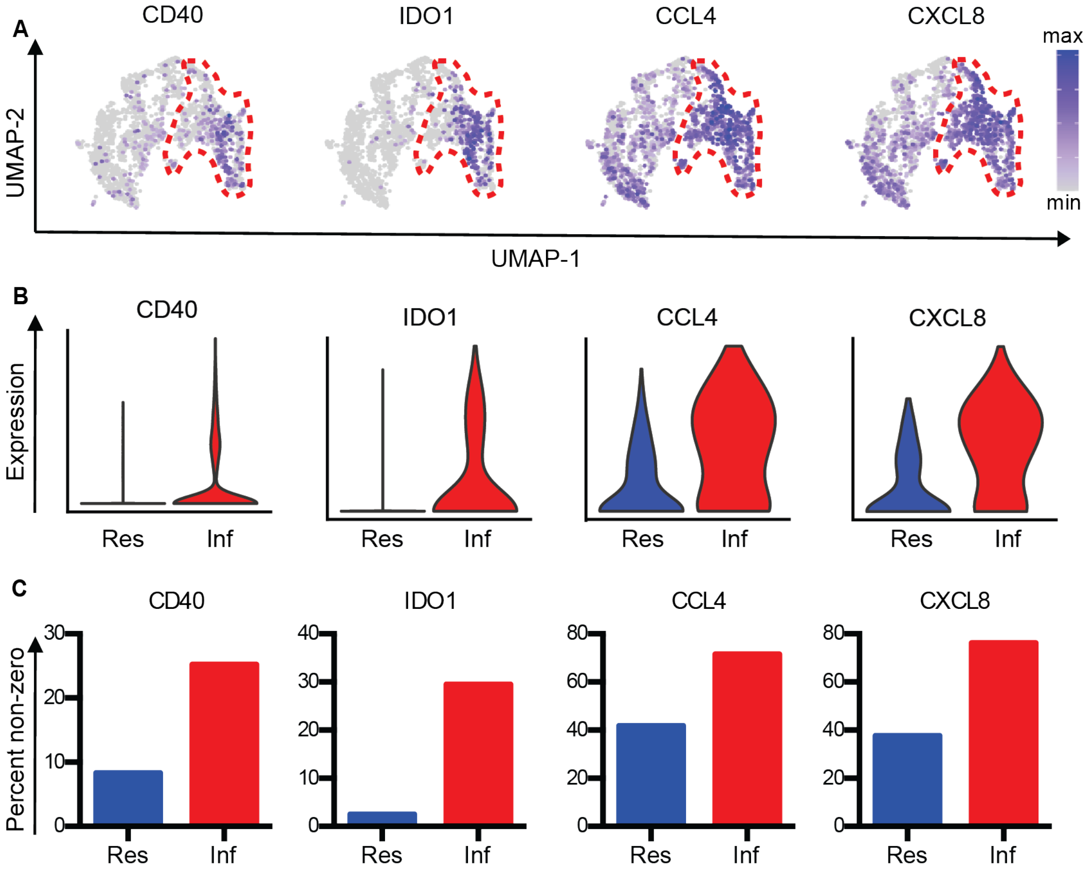


**Figure S11: Activation of IFN-induced and inflammatory pathways in macrophages from patients with inflammatory bowel disease.** A) UMAP plot from Fig. 5G showing gene expression in ileal macrophage populations. B) Violin plots of gene expression in macrophage clusters from Fig. 5G. C) Percent of cells with expression of genes in macrophage clusters from Fig. 5G. Res, resident macrophages; Inf, Inflammatory macrophages.

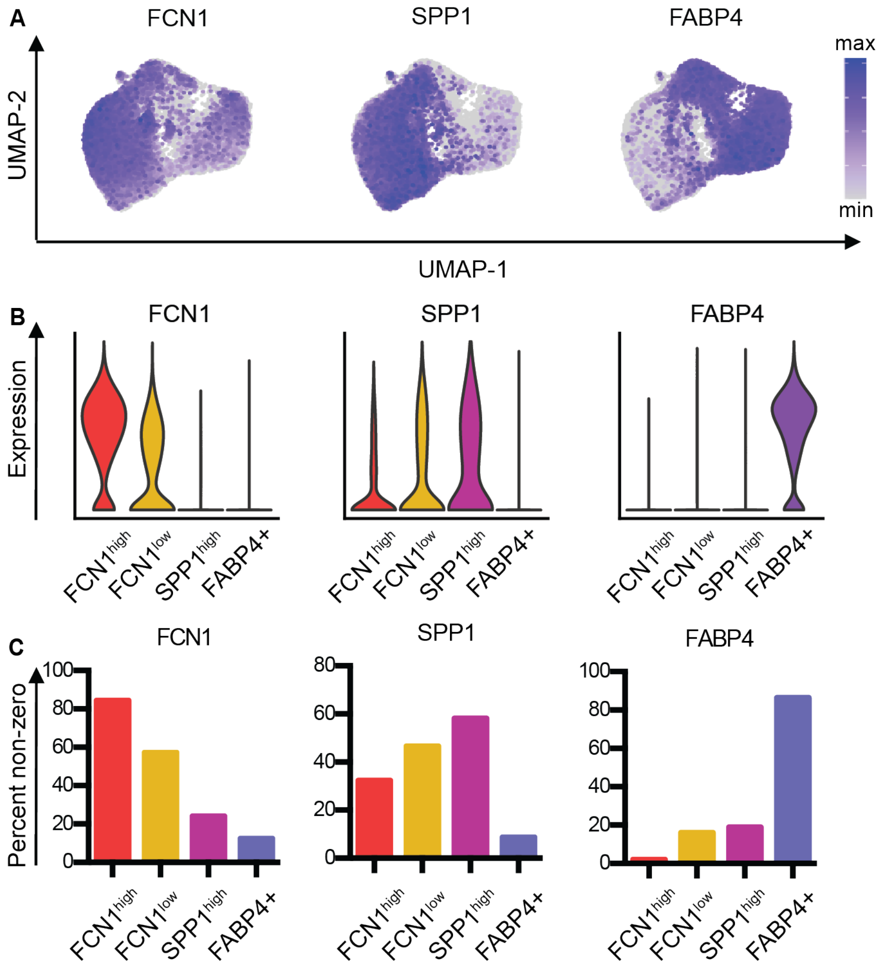


**Figure S12: Macrophage populations in bronchoalveolar lavage from patients with COVID-19.** A) UMAP plot from Fig. 5J showing gene expression in lung macrophage populations. B) Violin plots of gene expression in macrophage clusters from Fig. 5J. C) Percent of cells with expression of genes in macrophage clusters from Fig. 5J.

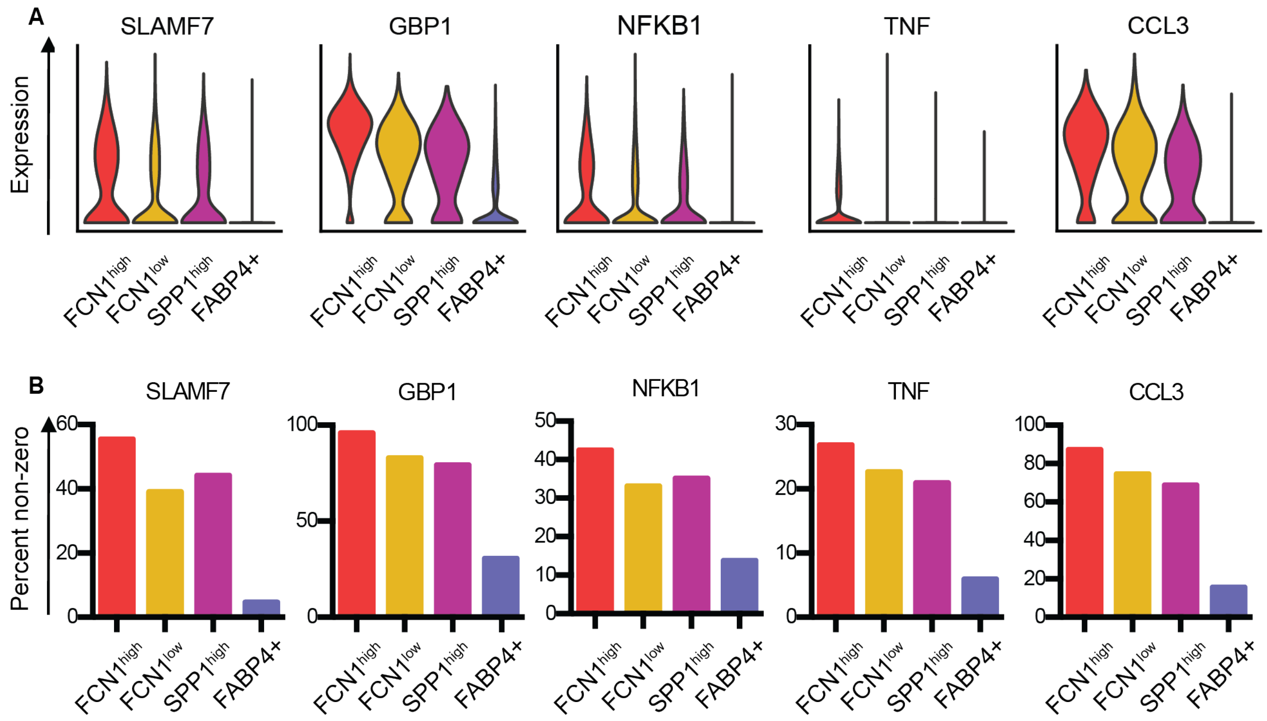


**Figure S13: Evidence for SLAMF7 activation in macrophages from patients with COVID-19 infection.** A) Violin plots of gene expression in macrophage clusters from Fig. 5J . B) Percent of cells with expression of genes in macrophage clusters from Fig. 5J.

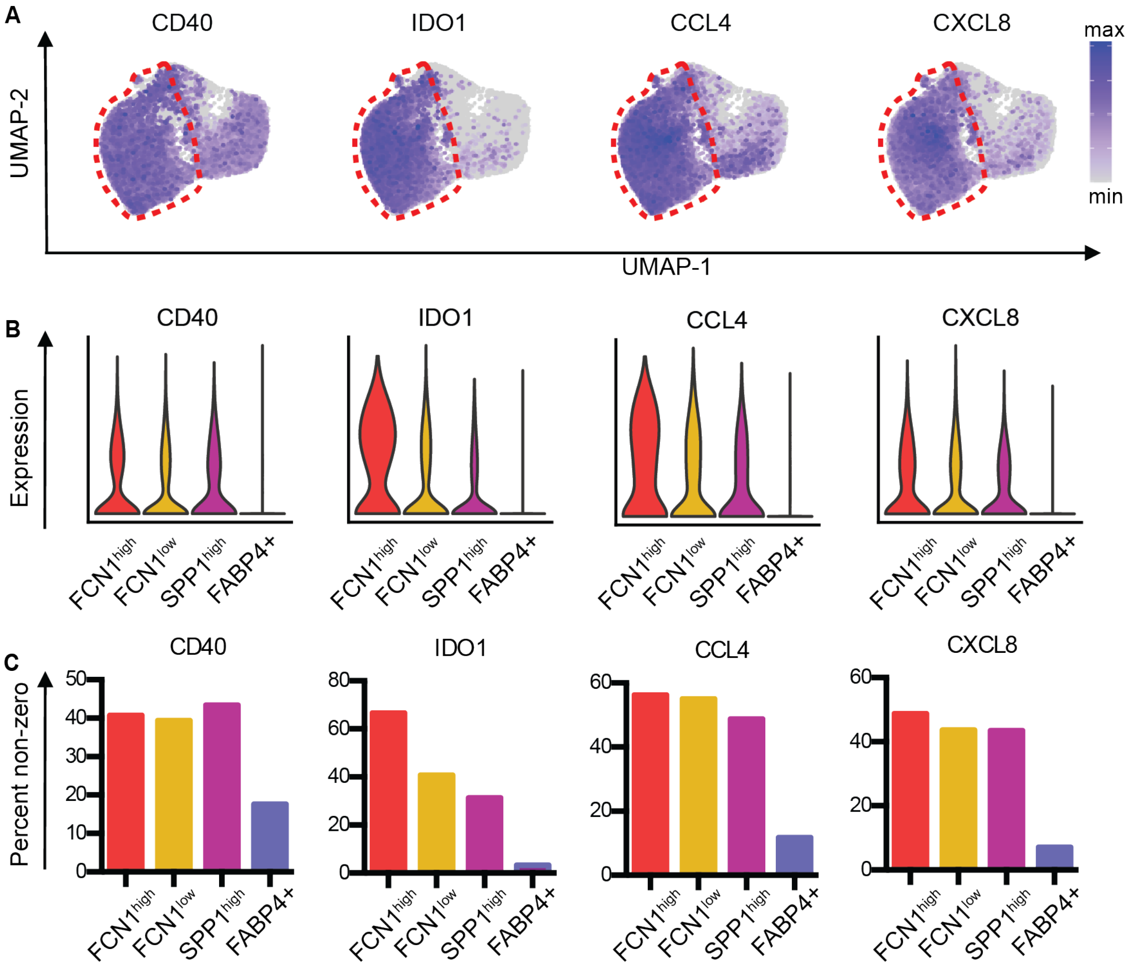


**Figure S14: Activation of IFN-induced and inflammatory pathways in macrophages from patients with COVID-19 infection.** A) UMAP plot from Fig. 5J showing gene expression in lung macrophage populations. B) Violin plots of gene expression in macrophage clusters from Fig. 5J. C) Percent of cells with expression of genes in macrophage clusters from Fig. 5J.
